# Body temperature maintenance acclimates in a winter-tenacious songbird

**DOI:** 10.1101/2020.03.27.012096

**Authors:** Maria Stager, Nathan R. Senner, Bret W. Tobalske, Zachary A. Cheviron

## Abstract

Flexibility in heat generation and dissipation mechanisms provides endotherms the ability to match their thermoregulatory strategy with external demands. However, the degree to which these two mechanisms account for seasonal changes in body temperature regulation is unexplored. Here we present novel data on the regulation of avian body temperature to investigate how birds alter mechanisms of heat production and heat conservation to deal with variation in ambient conditions. We subjected Dark-eyed Juncos (*Junco hyemalis*) to chronic cold acclimations of varying duration and subsequently quantified their metabolic rates, thermal conductance, and ability to maintain normothermia. Cold-acclimated birds adjusted traits related to both heat generation (increased summit metabolic rate) and heat conservation (decreased conductance) to improve their body temperature regulation. Increases in summit metabolic rate occurred rapidly, but plateaued after one week of cold exposure. In contrast, changes to conductance occurred only after nine weeks of cold exposure. Thus, the ability to maintain body temperature continued to improve throughout the experiment, but the mechanisms underlying this improvement changed through time. Our results demonstrate the ability of birds to adjust thermoregulatory strategies in response to thermal cues and reveal that birds may combine multiple responses to meet the specific demands of their environments.

## INTRODUCTION

Body temperature (T_b_) influences all aspects of animal function, from the rate of chemical reactions to metabolism, growth, and locomotion. Endogenous heat generation allows homeothermic endotherms to maintain a relatively constant T_b_ across a broad range of environmental temperatures, thereby providing physiological advantages (Bennett and Ruben, 1979; Crompton et al., 1978) that have enabled them to occupy a wide variety of habitats and climates. To maintain this high internal temperature, homeothermic endotherms coordinate changes occurring at multiple hierarchical levels of biological organization to respond to fluctuations in their environment.

The demands of T_b_ regulation are especially pronounced in temperate biomes where climates are often cooler than endothermic T_b_. Winter, in particular, can impose large temperature differentials for resident endotherms, and this thermoregulatory challenge is layered on top of other stresses, including reduced food availability, decreased daylight for foraging, and long nights of fasting (Marsh and Dawson, 1989). Unlike mammals that hibernate, a wide variety of birds remain active in temperate biomes all winter (Swanson, 2010). Some birds make use of heat-conservation mechanisms to cope with these conditions, such as huddling and utilizing microclimatic refugia, or employ facultative heterothermia, thereby decreasing their temperature differential with the environment and reducing energy consumption (Douglas et al., 2017; Korhonen, 1981; Mckechnie and Lovegrove, 2002). In spite of the benefits of these mechanisms, birds still need to eat, and they can frequently be seen foraging on even the most blustery days.

To remain active throughout the temperate winter, birds employ two primary physiological strategies to achieve normothermia. First, they can increase heat production. In general, avian thermogenesis results from shivering (Marsh and Dawson, 1989) or as a by-product of metabolism and activity (Dawson and O’Connor, 1996), although the role of non-shivering thermogenesis in adult birds is not well characterized (Hohtola, 2002). Peak oxygen consumption under cold exposure (summit metabolic rate; M_sum_) is often used as a proxy for thermogenic capacity, and many birds have been shown to increase M_sum_ by 10-50% in winter (Swanson, 2010). These seasonal changes have been credited with higher heat production and increased cold tolerance (O’Connor, 1995; Swanson, 1990a). At the same time, fueling an elevated metabolic rate requires increased foraging— and thus concomitantly escalates exposure to predators (Lima, 1985)—in addition to the potential energetic cost of restructuring internal physiology to meet these heightened aerobic demands (Liknes and Swanson, 2011). Few studies, however, have fully explored these potential trade-offs in natural systems (but see Petit et al., 2017), and shivering thermogenesis is frequently thought to represent the major mechanism by which birds maintain normothermia in winter (Swanson, 2010). Nonetheless, improved cold tolerance can occur independent of increases in M_sum_ (Dawson and Smith, 1986; Saarela et al., 1989), indicating additional strategies may be employed.

For small passerines that have high surface to volume ratios, seasonal decreases in thermal conductance (*i.e.* the transport of energy across a temperature gradient) may also be favored by natural selection. Direct measures of heat transfer are scarce (Wolf and Walsberg, 2000), but indirect measures indicate that thermal conductance decreases with decreasing ambient temperature across avian taxa (Londoño et al., 2017), which may be associated with increases in plumage density (Osváth et al., 2018). However, the role of seasonal adjustments to thermal conductance in birds is not well understood. Although some birds increase plumage mass in winter (Møller, 2015), it is unclear how this is achieved: most passerines molt only once per year, and their winter feathers are thus also their eventual summer feathers. Birds could also make behavioral adjustments in the cold, including postural changes to reduce surface area—especially of unfeathered areas, like the head and feet (Ferretti et al., 2019)—or erecting feathers to trap air around the body (Morris, 1956). Given these knowledge gaps, the question remains: what are the relative contributions of heat conservation and heat generation processes to avian body temperature regulation in the cold?

Such questions are particularly important in this era of rapid climatic change. Although ambient conditions can vary predictably, recent increases in climatic variability (*e.g.,* Kolstad et al., 2010) highlight the need for animals to respond rapidly to changing conditions. Each of the aforementioned potential physiological responses is likely tied to different environmental cues— primarily photoperiod and temperature (Swanson & Vézina, 2015). However, we do not understand how birds respond to environmental stimuli to balance heat loss and heat production, which is vital to projections of endothermic distributions under predicted future climate change scenarios (Buckley et al., 2018).

To understand how birds modify their thermoregulatory ability in the cold, we performed an acclimation experiment using Dark-eyed Juncos (*Junco hyemalis*). Juncos are small songbirds that overwinter across much of North America and are not known to huddle or use torpor (Nolan, Jr. et al., 2002). We exposed juncos sampled from a single population to one of ten experimental treatments that varied in temperature and the duration of cold exposure. Following acclimation to these experimental treatments, we quantified metabolic rates, heat loss across the skin and plumage, and T_b_ maintenance within the same individuals. Our results shed light on the ability of birds to respond to thermal cues and elucidate the mechanisms underlying their physiological responses to cold temperatures.

## MATERIALS AND METHODS

### Acclimation experiments

We captured adult juncos breeding in Missoula County, Montana, USA (~47.0°N, −113.4°W) from 12-19 July 2016 (*n* = 56) using mist nets. To increase sample sizes, we captured additional individuals 27 July - 3 August 2017 (*n* = 52) and repeated all procedures. We immediately transferred birds to husbandry facilities at the University of Montana and housed them individually under common conditions for 42 days (18°C, 10h light : 14h dark). After this six-week adjustment period, we assayed metabolic rates (see below). Following metabolic trials, we allowed birds to recover for ~24 hrs before we randomly assigned individuals to acclimation groups and subjected them to one of two temperature treatments, *Cold* (−8°C) or *Control* (18°C), lasting 7 d (*Week 1*), 14 d (*Week 2*), 21 d (*Week 3*), 42 d (*Week 6*), or 63 d (*Week 9*) in duration. We chose to acclimate birds to −8°C, which is a temperature that juncos experience in the northern parts of their winter range for weeks at a time (Fig. S1) and which could elicit more dramatic physiological responses than previous experiments with juncos performed at 3°C (Swanson et al., 2014). Photoperiod was maintained at a constant 10L: 14D in both treatments (the photoperiod in Missoula County in November and February), and food and water were supplied *ad libitum* for the duration of the experiment. Birds were fed white millet and black oil sunflower seeds at a 2:1 ratio by weight, supplemented with ground dog food, live mealworms, and vitamin drops (Wild Harvest D13123 Multi Drops) in their water. We did not repeat the *Week 9* treatment in 2017. Also, one individual died 12 days into the *Cold* treatment in 2016 and another died during the adjustment period in 2017 (causes unknown), resulting in a total sample size of 106 individuals (*n* = 12 per treatment, except *n_Control_1_* =11, *n_Control_9_ = 6*, *n_Cold_9_* = 5).

As an index of body size, we measured the tarsus lengths (mm) of both legs and calculated the average measure for each individual. We quantified this feature only once (after the bird was euthanized) assuming that tarsus length did not change over the duration of the acclimation because all individuals were adults. The sample is heavily male-biased (90.5%) but includes 10 females (9.5%) across the two years. These females were randomly distributed across most treatment groups (Table S1). Brood patches and cloacal protuberances were not present after the six-week adjustment period. Sex was confirmed post-acclimation by identification of the gonads during dissection. For five additional males captured at the same time but not included in the study, we confirmed by dissection that testes had regressed before the acclimations began.

### Metabolic assays

We measured resting metabolic rate (RMR) and M_sum_ in a temperature-controlled cabinet using open-flow respirometry before and after acclimation treatments. RMR trials were conducted in the evening during birds’ dark cycle (start time *μ* = 19:11; range = 18:00 – 23:20). M_sum_ trials were conducted the following day largely within birds’ light cycle (start time *μ* = 13:30; range = 09:00 – 20:42). Birds were not fasted before either measurement so as not to limit aerobic performance and to ease comparison between measures. For RMR trials, birds were placed in a modified 1-L Nalgene container and measured in a dark, quiet temperature cabinet (Sable Systems Pelt Cabinet with Pelt-5 Temperature Controller) at 27° C, which is within the thermoneutral zone of juncos (Swanson, 1991). Three individuals were assayed simultaneously. We cycled through individuals at 15-min intervals alternated with 5-min ambient baseline measures, such that each individual was measured for at least 30 minutes over the course of 2 hours. We subjected an individual to additional rounds of measurement if the O_2_ trace suggested that it was active. Ambient air was first dried (using drierite), pumped through the animal chamber at 500 ml/min, and excurrent air was subsampled manually from one chamber at a time at 100–150 ml/min through barrel syringes. We dried excurrent air again, then CO_2_ was scrubbed with ascarite, and the outflow dried again before passing through a FoxBox (Sable Systems) to quantify O_2_. The same setup was used for both baseline and animal chambers. We spanned the FoxBox using baseline air at 20.95% O_2_ before each trial began. Flow was controlled using a mass flow meter (Sable Systems). From these measures, we quantified oxygen consumption according to Lighton (2008). We first corrected for any fluctuations in baseline concentrations then calculated RMR as the lowest oxygen consumption (ml O_2_•min^−1^) averaged over a ten-minute period using custom scripts in the R programming environment (R Core Team, 2018).

M_sum_ trials were conducted using a similar setup with static cold exposure. Trials were conducted in a heliox environment (21% helium, 79% oxygen) with flow rates of 750 ml/min. The high thermal conductance of heliox facilitates heat loss at higher temperatures than is necessary in air to avoid injury to experimental subjects (Rosenmann and Morrison, 1974). Heliox flow rates were measured using a mass flow meter (Alicat M-series) programmed for the specific gas mixture. Pre-acclimation M_sum_ trials were conducted using the above temperature cabinet set to −5° C. Trials ended when a bird‘s CO_2_ production plateaued or after one hour, whichever came first. Immediately upon removing birds from the temperature cabinet, we measured body temperature using a thermistor probe inserted into the cloaca. We considered birds hypothermic if their body temperature was ≤ 37° C (*per* Swanson et al., 2014). One individual that was not hypothermic at the end of the M_sum_ trail was removed from further analysis. We corrected for drift then calculated M_sum_ as the highest oxygen consumption (ml O_2_•min^−1^) averaged over a five-minute period using custom scripts in R. As a measure of thermogenic endurance, we calculated the number of minutes that an individual maintained 90% or more of their M_sum_ (Cheviron et al., 2013).

Because we expected acclimated birds to differ in their cold tolerance, we performed post-acclimation M_sum_ trials at lower temperatures for cold acclimated birds (starting cabinet temperature *μ* = − 24.47 ± 2.87) than control acclimated birds (*μ* = −15.94 ± 5.98) using a laboratory freezer (Accucold VLT650). These temperatures, concurrent with a heliox atmosphere, represent rather severe conditions that juncos are unlikely to encounter in the wild but were chosen because previous work has demonstrated that cold exposure in excess of −9° C in heliox is necessary to induce hypothermia within 90 minutes in winter acclimatized juncos (Swanson, 1990a). Although we aimed for static cold exposure, logistical constraints did not allow for precise temperature control. We thus recorded temperature inside the cabinet for the duration of the trial to account for variation within and among trials. Post-acclimation trials ended after an extended period of declining CO_2_ production coincident with the bird’s body temperature dropping below 30° C (see below).

We used multiple respirometry setups in order to complete all pre-acclimation measurements precisely 42 d after the day of capture (three units in 2016, four in 2017). Post-hoc tests revealed significant differences in the metabolic measurements made by each respirometry unit. To control for these effects, we regressed each metabolic trait (RMR or M_sum_) on respirometry unit for each year and then subtracted the resulting beta coefficient (slope) from the metabolic rate (Table S2). Although all post-acclimation measures were conducted using a single respirometer, we used the same correction factor to make the before and after measures comparable. In a few instances, this resulted in negative M_sum_ values that were removed from further analysis (*n* = 3 pre-acclimation measures, *n* = 1 post-acclimation). Metabolic trials for cold individuals were conducted earlier in the day than those of control individuals because the temperature cabinet tended to increase in temperature each time it was opened. For this reason, we tested for but did not find a significant interaction between trial start time and temperature treatment on post-acclimation M_sum_ (*p* = 0.21).

We measured body mass (M_b_; in g) immediately before each metabolic assay. Birds were banded with a unique combination of two or three plastic leg bands; the mass of these bands has been removed from all reported M_b_. Directly following the post-acclimation M_sum_ trial, we euthanized individuals using cervical dislocation, removed organs and tissues within the body cavity, filled the body cavity with a wet paper towel to preserve moisture, and froze carcasses at −20° C until thermal conductance assays were performed in May and June 2019. To quantify the change in each trait value with acclimation, we subtracted an individual’s pre-acclimation trait value from their post-acclimation value (ΔM_b_, ΔRMR, and ΔM_sum_). We did not compare endurance measures pre- and post-acclimation because trial conditions varied before and after acclimation.

### Thermal conductance assays

We measured the conductive properties of the skin and plumage by quantifying the amount of power input (mW) required to maintain a constant internal temperature of 39°C with the ambient temperature providing a gradient. To do this, we first thawed carcasses at room temperature and dried the feathers. We removed any adipose or muscle tissue remaining in the body cavity, then inserted an epoxy mold (~35 mm long x 16 mm in diameter; PC-Marine Epoxy Putty) into the coelom that we designed to fill the coelom without significant stretching of the superficial thoracic and abdominal regions. Within this mold, we embedded a centrally placed thermocouple and a length of nichrome wire for heating. These were connected to a custom-made board containing a Voltage logger (Omega OM-CP-Quadvolt), an amperage logger (Omega OM-CP-Process 101A-3A), and a temperature controller (Omega CNI1622-C24-DC). Power was supplied to the circuit using a 12V DC battery. We sewed the body cavity together using sewing thread, leaving a small hole near the cloaca for the wires to exit. We suspended the carcass from a single thread through the nares, supported by the wires from below, such that birds were in an upright position with legs hanging freely. We cleaned the feathers with cornmeal to remove oils and combed the feathers into place. Wings were positioned at the sides, tucked in as best as possible. We removed 6 carcasses damaged beyond repair in post-processing.

Conductance trials were conducted in a small, closed room without airflow and at ambient (laboratory) temperature (*μ* = 23.4 ± 0.61). The mold was first brought to 39° C and power was supplied whenever the temperature dropped below 38° C. We recorded the amperage, Voltage, and temperature of the thermocouple for each second of an eighteen-minute trial. We calculated the average power input (conductance, mW) as the mean Volts × amps over a ten-minute period. We excluded two individuals for which temperatures did not stay within the specified range, resulting in a total sample size of *n* = 98. All assays were performed by a single individual (MS) and were done blind to the birds’ treatment assignments. We did not find a significant effect of the minor variation in ambient temperature that occurred on average power input using a linear regression (*p* = 0.19). Trials were performed across multiple days, but we did not find an effect of measurement day (Table S3) or freeze duration on average power input (*p* = 0.95).

### Body temperature maintenance

To quantify the ability to maintain normothermia during acute cold exposure, we measured T_b_ continuously for the duration of the post-acclimation M_sum_ acute cold trial. Immediately prior to this trial, we inserted a temperature-sensitive passive integrated transponder (PIT) tag (12mm, Biomark) into the cloaca of the bird. PIT tags were inserted at room temperature; thus, even *Cold* birds were exposed to warmer conditions for a few minutes preceding the M_sum_ trial. To secure the tag, we glued the feathers surrounding the cloaca together using cyanoacrylate adhesive (super glue). We quantified M_b_ before the addition of the PIT tag. An antenna was placed inside the temperature cabinet next to the animal chamber and connected to an external reader that recorded T_b_ eight times per second (Biomark HPR Plus Reader). We averaged the T_b_ measurements over each one-minute interval of the trial and coded each one-minute interval as hypothermic or normothermic. We deemed birds hypothermic once they lost 10% of their initial T_b_ and maintained T_b_ below this level. Because birds differed in their initial T_b_ (36-42° C), we repeated all analyses using the commonly accepted threshold of 37.0° C to define the hypothermic state, but this did not change our overall results (Tables S4-S5). In some cases, super glue did not hold the cloaca closed, and birds ejected their PIT tags during the trial. We removed from the sample 6 individuals for which PIT tag ejection occurred before hypothermia could be assessed. We also removed 8 individuals for which gaps longer than one minute existed (due to the position of the bird relative to the antenna) at critical periods that prevented precise detection of their hypothermic state, resulting in a total sample size of *n* = 92. We used different respirometry chambers (either a custom-made plexiglass box or modified Nalgene) for the post-acclimation M_sum_ trials between years. Because these chambers had different thermal properties that may have contributed to differences in the way the individuals experienced temperature in the cold trials, we also tested for an effect of Year on risk of hypothermia (see below).

### Analyses

We performed all analyses in R. We first quantified the effects of acclimation temperature and duration on mass, tarsus length, and conductance using multiple regressions for pre-acclimation, post-acclimation, and ΔM_b_ values. We similarly used multiple regressions to quantify the effects of acclimation temperature and duration on RMR, M_sum_, and endurance with M_b_ as a covariate, as well as on ΔRMR and ΔM_sum_ with ΔM_b_ as a covariate. For all models, we also tested for an effect of a temperature × duration interaction but this term was generally not significant (Table S6). Additionally, we tested for associations among the phenotypic traits using Pearson correlation tests. We report means ± standard deviations in the text.

To assess T_b_ maintenance, we used T_b_ interval data to fit Cox proportional hazards regression models using the *survival* package in R (Therneau, 2015). These standard time to event models analyse non-linear processes without assuming any one shape of response, allowing us to control for differences in temperature stimulus among individuals. We created survival objects with interval data and hypothermic status, then fit regressions using the function *coxph* to quantify the effects of cabinet temperature, temperature treatment, duration, and year with all terms clustered by individual on the risk of hypothermia. We first standardized all variables using the *arm* package (Gelman, 2008).

We used the same approach to assess the effect of phenotypic traits (M_b_, tarsus, RMR, M_sum_, endurance, and conductance) on the risk of hypothermia using a subset of individuals for which we had complete measurements (*n* = 84). Because of the large number of phenotypic variables potentially influencing T_b_ maintenance, we used a model selection process whereby we tested all possible combinations (including two-way interactions) of the predictor variables. We evaluated all models using Akaike information criterion scores corrected for small sample sizes (AIC_c_), where the model with the lowest AIC_c_ score was considered the most well supported model. Because there was no single most well supported model (e.g., *w_i_* > 0.90; Grueber et al., 2011), we used model averaging to identify which predictor variables had significant effects on T_b_ maintenance.

## RESULTS

Prior to acclimation, treatment groups did not differ significantly in body size or metabolic traits (Table 1). Acclimation temperature and duration did not influence M_b_ (*μ* = 22.30 ± 1.79 g) or RMR (*μ* = 1.38 ± 0.29 ml O_2_•min^−1^; Table 1; Figure 1a). RMR was correlated with M_b_ both before and after acclimation (Table 1).

**Table 1.** Linear effects of *Cold* treatment and *Duration* on phenotypic traits before and after acclimation. Mass (M_b_) is included as a covariate for metabolic traits. Delta (Δ) represents change over acclimation period (post-minus pre-acclimation) for traits that were measured at both time points. Bolded significant effects after Bonferroni correction for multiple models (p < 0.004).

| | Phenotype | $n$ | Intercept | | $M_b$ | | | Cold Treatment | | | Duration | | | Adj. $R^2$ |
| --- | --- | --- | --- | --- | --- | --- | --- | --- | --- | --- | --- | --- | --- | --- |
| | | | $\beta$ | SE | $\beta$ | SE | $p$ | $\beta$ | SE | $p$ | $\beta$ | SE | $p$ | |
| Pre | $M_b$ | 106 | 22.34 | 0.31 | | | | 0.02 | 0.31 | 0.94 | -0.05 | 0.06 | 0.41 | -0.01 |
|  | Tarsus | 106 | 19.91 | 0.11 |  |  |  | 0.10 | 0.11 | 0.33 | 0.04 | 0.02 | 0.07 | 0.02 |
|  | **RMR** | 106 | 0.28 | 0.31 | **0.05** | **0.01** | **$8.9 \times 10^{-4}$** | -0.01 | 0.04 | 0.82 | -0.01 | 0.01 | 0.13 | 0.10 |
| | $M_{sum}$ | 102 | 4.90 | 0.79 | 0.06 | 0.04 | 0.07 | 0.10 | 0.12 | 0.40 | -0.01 | 0.02 | 0.52 | 0.02 |
|  | Endur. | 103 | 37.56 | 18.20 | -0.48 | 0.81 | 0.55 | 1.52 | 2.76 | 0.58 | 0.05 | 0.55 | 0.93 | -0.02 |
| Post | $M_b$ | 106 | 22.83 | 0.33 | | | | 0.47 | 0.32 | 0.14 | -0.13 | 0.06 | 0.04 | 0.04 |
|  | **RMR** | 105 | 0.30 | 0.43 | **0.05** | **0.02** | **$2.3 \times 10^{-3}$** | -0.04 | 0.05 | 0.51 | -0.02 | 0.01 | 0.13 | 0.10 |
| | $M_{sum}$ | 105 | 5.12 | 1.71 | 0.07 | 0.07 | 0.31 | **1.31** | **0.25** | **$8.1 \times 10^{-7}$** | -0.10 | 0.05 | 0.07 | 0.23 |
|  | Endur. | 105 | 17.88 | 21.85 | 0.15 | 0.93 | 0.87 | -6.04 | 3.19 | 0.42 | 1.24 | 0.65 | 0.06 | 0.04 |
|  | Conduct. | 98 | 323.63 | 6.89 |  |  |  | -0.50 | 6.89 | 0.94 | -1.07 | 1.35 | 0.43 | -0.01 |
| $\Delta$ | $M_b$ | 106 | 0.49 | 0.37 | | | | 0.45 | 0.37 | 0.22 | -0.08 | 0.07 | 0.28 | 0.01 |
|  | **RMR** | 106 | 0.10 | 0.06 | **0.05** | **0.02** | **$2.2 \times 10^{-3}$** | -0.02 | 0.06 | 0.71 | 0.00 | 0.00 | 0.82 | 0.06 |
| | $M_{sum}$ | 102 | 0.57 | 0.26 | -0.10 | 0.06 | 0.12 | **1.14** | **0.24** | **$1.2 \times 10^{-5}$** | -0.09 | 0.05 | 0.06 | 0.19 |

**Figure 1.**
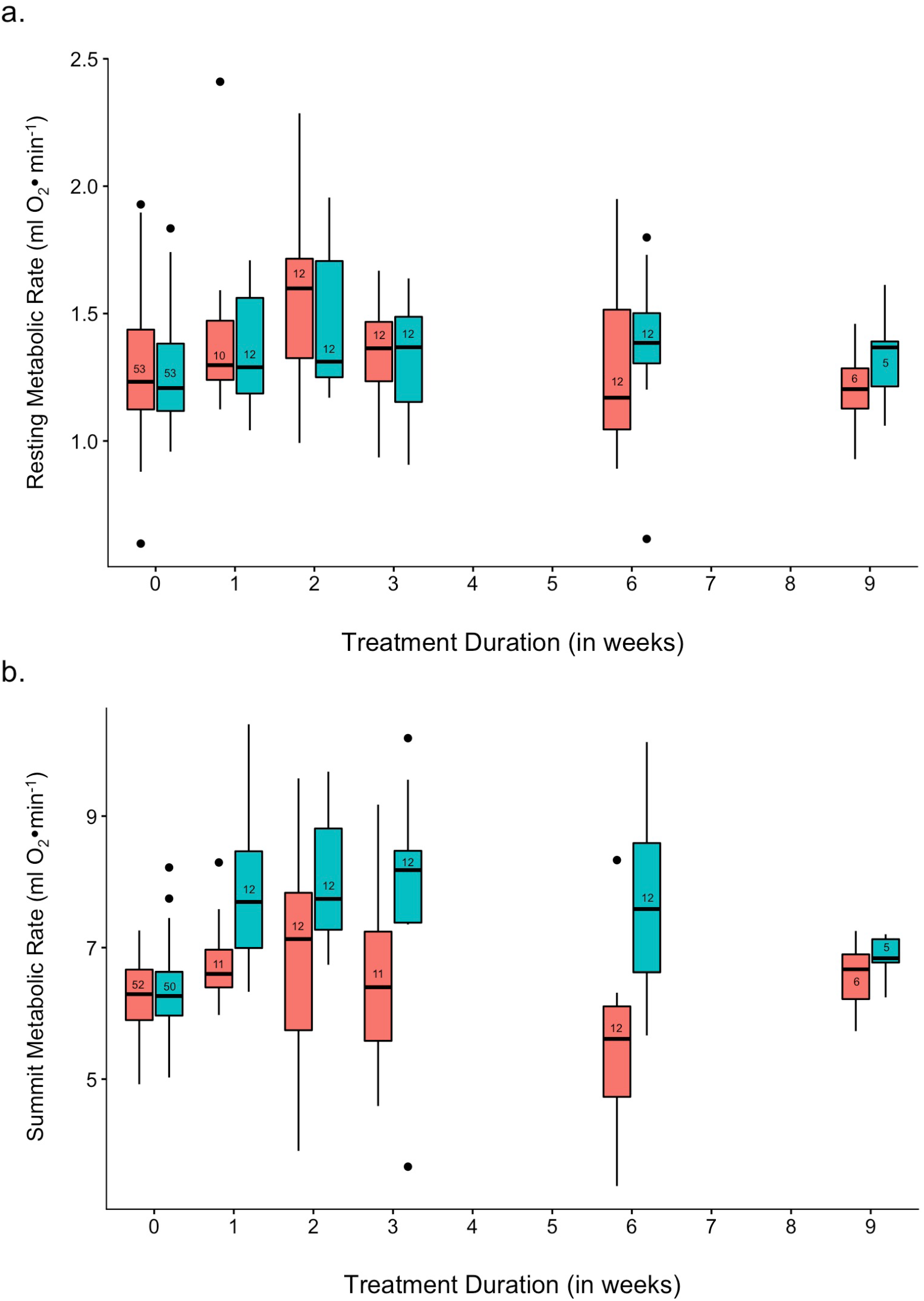
(a) Resting and (b) summit metabolic rate of juncos across treatments. Pre-acclimation measures for all individuals shown at Week 0. Numbers in boxes indicate sample sizes for each group. Red = *Control*; Blue = *Cold*. Boxplots show the median values (horizontal line in the box), the 25th and 75th percentiles (lower and upper margins of the box) together with the minimum and maximum values ≤ 1.5 * IQR from the box margin (whiskers), and outlying points (circles).

In contrast, cold-acclimated birds exhibited a 20% elevation in M_sum_ compared to *Control* birds (Table 1; Figure 1b). Duration of cold-exposure did not influence M_sum_ and M_sum_ was not correlated with M_b_ before or after acclimation (Table 1). Similarly, M_sum_ did not correlate with RMR at either time point (*r_pre_* = −0.01, *p_pre_* = 0.85; *r_post_* = 0.15, *p_post_* = 0.13). Thermogenic endurance did not vary with temperature treatment or duration (Table 1), nor did it correlate with M_sum_ (*r* = −0.16, *p* = 0.11).

Conductance properties of the skin were largely unchanged across acclimation treatments (Table 1). However, there was an interaction between treatment and duration (*β* = −8.52 ± 2.56, *p* = 0.0013). To investigate this relationship, we reran our regression model with Duration as a categorical rather than continuous variable (Table 2). This revealed that the skin and plumage of *Cold Week 9* birds exhibited a reduction in heat transfer compared to other groups (Figure 2). The average power input required to maintain core temperature at 39° C was not correlated with M_b_ (*r* = −0.14, *p* = 0.16), tarsus length (*r* = −0.11, *p* = 0.29), RMR (*r* = −0.14, *p* = 0.16), or M_sum_ (*r* = 0.03, *p* = 0.77).

**Table 2.** Linear effects of *Treatment*, *Duration* (as categorical variable), and their interaction on conductance properties of the skin and plumage. *Control Week 1* is reference.

| Variable | $\beta$ | SE | $p$ |
| --- | --- | --- | --- |
| **Intercept** | 316.88 | 10.06 | $<2.0 \times 10^{-16}$ |
| **Treatment** | 16.37 | 13.91 | 0.24 |
| **Week 2** | -15.51 | 13.62 | 0.26 |
| **Week 3** | 12.15 | 13.63 | 0.37 |
| **Week 6** | 6.31 | 14.62 | 0.67 |
| **Week 9** | 20.55 | 16.44 | 0.21 |
| **Treatment × Week 2** | -4.38 | 19.23 | 0.82 |
| **Treatment × Week 3** | -9.65 | 19.03 | 0.61 |
| **Treatment × Week 6** | -28.78 | 20.18 | 0.16 |
| Treatment × Week 9 | **-74.36** | **23.77** | **$2.4 \times 10^{-3}$** |

**Figure 2.**
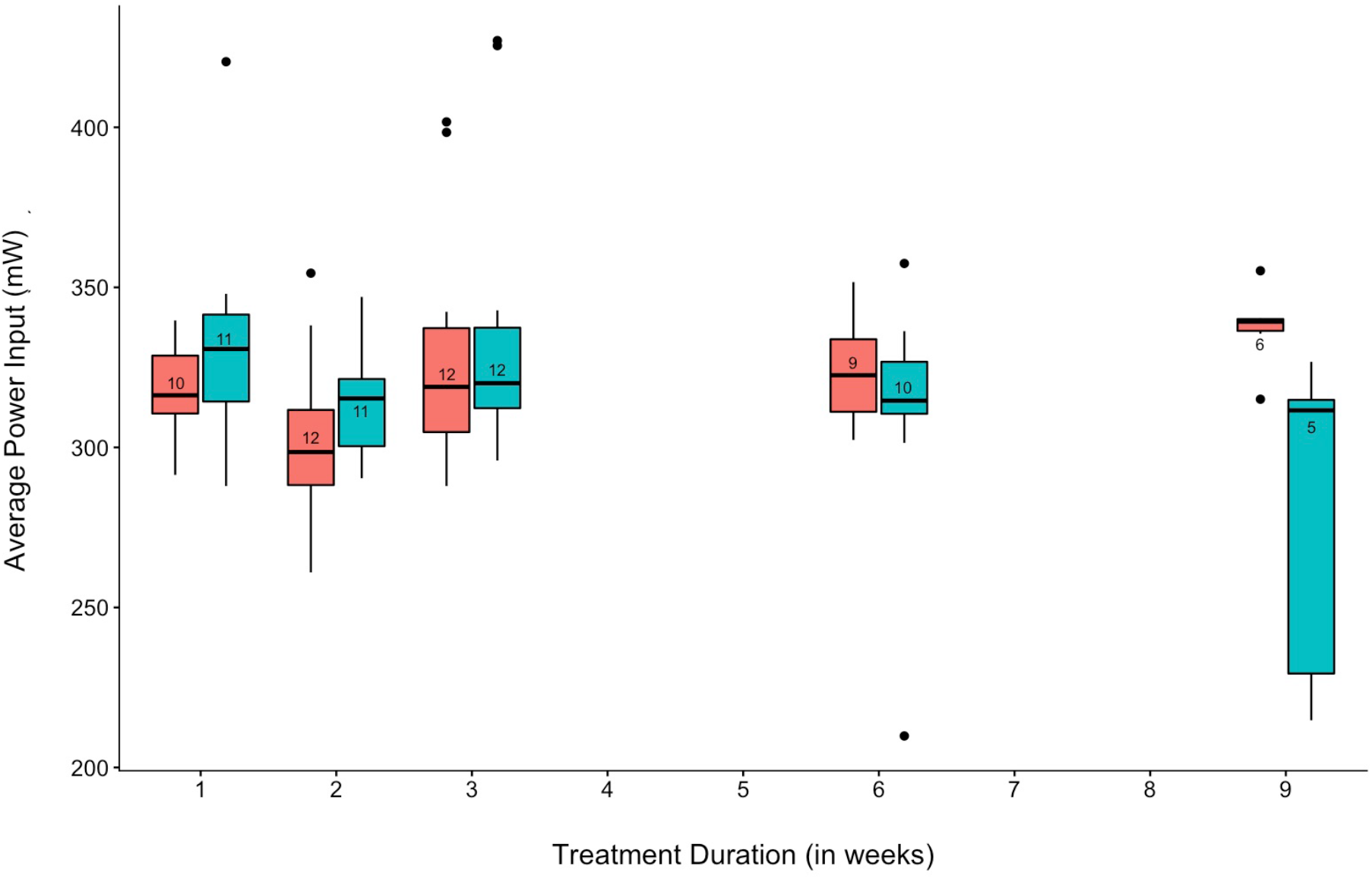
Heat loss properties of junco skins across treatment groups expressed as the power (mW) required to maintain core body temperature at 39°C with ambient temperature at 24°C. Numbers in boxes indicate sample sizes for each group. Red = *Control*; Blue = *Cold*. For boxplot conventions, see legend to Fig. 1.

Temperature loss trajectories varied among individuals in acute cold trials. Some juncos showed a steady decline in T_b_ over time, while others exhibited an oscillating T_b_ (Figure 3). Thirteen individuals, distributed across treatment groups, demonstrated the ability to increase T_b_ above normothermia after sustaining substantial losses in T_b_. Birds did not differ in T_b_ among temperature acclimation groups at the start of the trial (t-test: t(94)=0.45, *p* = 0.65).

**Figure 3.**
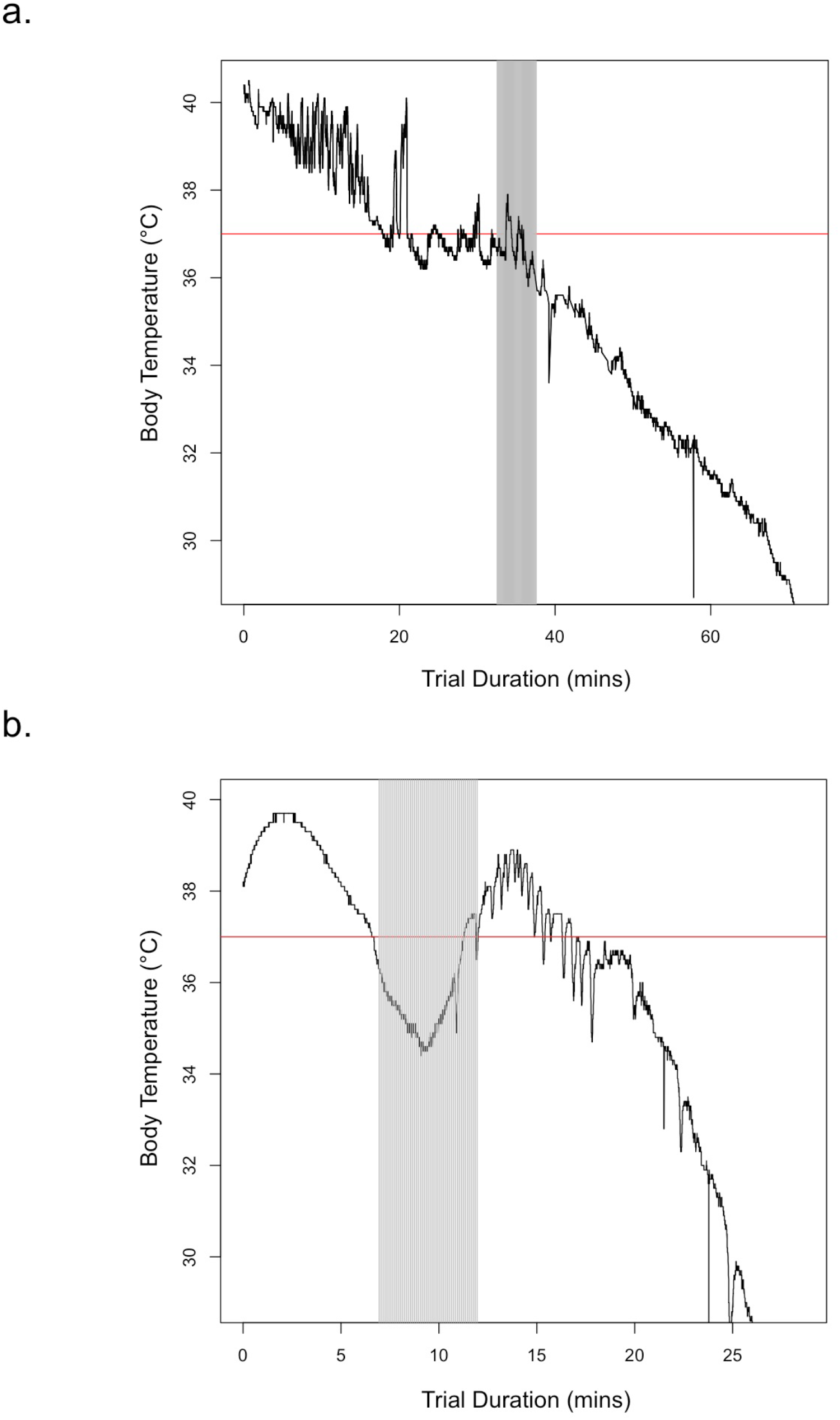
Example trajectories of body temperature loss during acute cold trials for an individual that exhibits (a) mostly continual loss and (b) one that regains normothermia. Black line = body temperature; red line = 37°C; gray box is the 5-minute period corresponding to M_sum_.

Higher cabinet temperatures elicited a reduced risk of hypothermia with a 17% reduction in per minute hazard for every 1° increase in cabinet temperature (Table 3). For this reason, we included cabinet temperature as a covariate in all subsequent models. Cold-acclimated birds exhibited an 87% reduction in the per minute risk of hypothermia in acute cold trials (Figure 4a). Every week of acclimation duration was associated with a 15% reduction in the per minute risk of hypothermia. This was true for both the *Cold* and the *Control* treatments, so to further investigate this relationship, we tested for the effect of duration as a categorical, rather than continuous variable. Within the *Control* treatment, only *Week 9* individuals showed a reduction in hypothermia risk compared to *Week 1* birds (Table 4a). However, within the cold-acclimated birds, *Weeks 2*, *6*, and *9* all showed a reduced risk of hypothermia compared to *Week 1* (Table 4b; Figure 4b). Year did not influence the risk of hypothermia (Table 3).

**Table 3.** Cox proportional hazards model output for T_b_ maintenance as a function of cabinet temperature, acclimation temperature treatment, duration treatment, and year. Negative *β* coefficients represent reduced risk of hypothermia. Hazards ratio (HR) is the exponent of the *β* coefficient (*i.e.* a reduction in the hazard by this factor). *Control* treatment is reference for temperature effect. All continuous variables were standardized; bold indicates predictor variables with statistically significant effects on T_b_ maintenance.

| Variable | $\beta$ | SE | HR | 95% CI | $p$ |
| --- | --- | --- | --- | --- | --- |
| **Cabinet Temp.** | **-3.87** | **0.55** | **0.02** | **-5.17, -2.58** | **$4.8 \times 10^{-9}$** |
| **Treatment** | **-2.06** | **0.48** | **0.13** | **-3.22, -0.90** | **$4.9 \times 10^{-4}$** |
| **Period** | **-0.95** | **0.26** | **0.39** | **-1.65, -0.26** | **$7.4 \times 10^{-3}$** |
| Year | 0.53 | 0.25 | 1.70 | -0.22, 1.28 | 0.17 |

**Table 4.** Survival model output for hypothermic state as a function of treatment group for (a) *Control* birds only and (b) *Cold* birds only. *Week 1* as reference. Negative *β* coefficients represent reduced risk of hypothermia. Hazards ratio (HR) is the exponent of the *β* coefficient (*i.e.* a reduction in the hazard by this factor). Bold indicates predictor variables with statistically significant effects on T_b_ maintenance.

| Variable | $\beta$ | SE | HR | $p$ |
| --- | --- | --- | --- | --- |
| Cabinet Temp. | **-0.15** | **0.06** | **0.86** | **0.05** |
| **Week 2** | -0.98 | 0.51 | 0.38 | 0.36 |
| **Week 3** | -1.27 | 0.50 | 0.28 | 0.14 |
| **Week 6** | 0.14 | 0.50 | 1.15 | 0.84 |
| **Week 9** | **-2.23** | **0.59** | **0.11** | **$1.8 \times 10^{-4}$** |

(b)
| Variable | $\beta$ | SE | HR | $p$ |
| --- | --- | --- | --- | --- |
| Cabinet Temp. | **-0.50** | **0.08** | **0.61** | **$2.5 \times 10^{-7}$** |
| Week 2 | **-1.82** | **0.52** | **0.16** | **$1.1 \times 10^{-3}$** |
| **Week 3** | -1.01 | 0.51 | 0.36 | 0.11 |
| Week 6 | **-1.23** | **0.49** | **0.29** | **0.01** |
| **Week 9** | **-4.44** | **0.84** | **0.01** | **$6.1 \times 10^{-4}$** |

**Figure 4.**
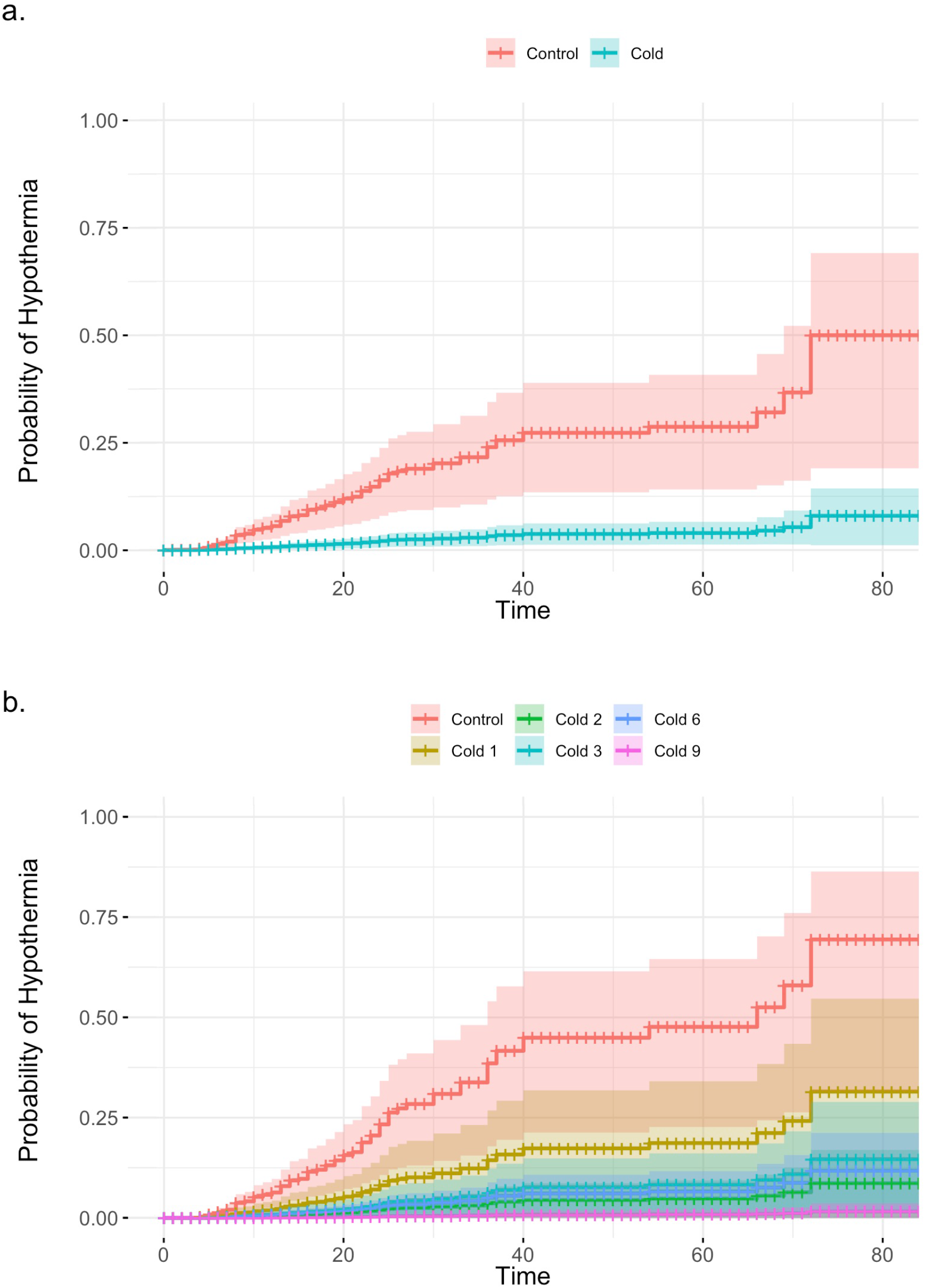
Survival curves depicting time to hypothermia in acute cold trials across (a) temperature (*n* = 92) and (b) duration treatments (*n* = 86) while controlling for cabinet temperature. *Control* treatments (excluding *Week 9*) combined in (b). Regression lines shown with shaded areas representing 95% confidence intervals.

There was no single model best predicting risk of hypothermia using phenotypic traits (Table 5). However, model averaging identified M_sum_, Endurance, and the interaction between M_sum_ × Endurance as significant predictor variables (Table 6). The interaction term indicates that birds with both higher M_sum_ and Endurance were better able to maintain their T_b_. In comparison, M_b_, tarsus length, RMR, and conductance were not correlated with time to hypothermia (Table 6).

**Table 5.** Highest-ranked models (with lowest AIC_c_ scores) in candidate set for effects of phenotypic variables on the maintenance of T_b_ using Cox proportional hazards models. Only models with ΔAIC_c_ < 4 are reported. *K* indicates the number of parameters in each model; Cabinet refers to the cabinet temperature during the cold trial.

| Candidate Model | *K* | AIC<sub>c</sub> | $\Delta\text{AIC}_c$ | $w_i$ |
| --- | --- | --- | --- | --- |
| **Cabinet + Endurance × M<sub>sum</sub> + RMR** | 5 | 1007.9 | 0.0 | 0.23 |
| **Cabinet + Conductance + Endurance × M<sub>sum</sub> + RMR** | 6 | 1009.2 | 1.3 | 0.12 |
| **Cabinet + Endurance × M<sub>sum</sub> + RMR + Tarsus** | 6 | 1009.6 | 1.7 | 0.10 |
| **Cabinet + Endurance × M<sub>sum</sub> + M<sub>b</sub> + RMR** | 6 | 1009.6 | 1.7 | 0.10 |
| **Cabinet + Conductance + Endurance × M<sub>sum</sub> + M<sub>b</sub> + RMR** | 7 | 1010.9 | 3.0 | 0.05 |
| **Cabinet + Endurance × M<sub>sum</sub> + M<sub>b</sub> + RMR + Tarsus** | 7 | 1010.9 | 3.0 | 0.05 |
| **Cabinet + Conductance + Endurance × M<sub>sum</sub> + RMR + Tarsus** | 7 | 1011.0 | 3.1 | 0.05 |
| **Cabinet + Endurance × M<sub>sum</sub>** | 4 | 1011.2 | 3.3 | 0.04 |
| **Cabinet + Conductance + Endurance × M<sub>sum</sub>** | 5 | 1011.3 | 3.4 | 0.04 |

**Table 6.** Model-averaged coefficients for phenotypic variables affecting the maintenance of T_b_ assessed using Cox proportional hazards models. Negative *β* coefficients represent reduced risk of hypothermia. Hazards ratio (HR) is the exponent of the *β* coefficient (*i.e.* a reduction in the hazard by this factor). All continuous variables were standardized; bold indicates predictor variables with statistically significant effects on T_b_ maintenance.

| Variable | $\beta$ | SE | HR | 95% CI | $p$ |
| --- | --- | --- | --- | --- | --- |
| Cabinet Temp. | -0.93 | 0.49 | 0.39 | -1.90, 0.02 | 0.06 |
| **Endurance** | **-2.02** | **0.61** | **0.13** | **-3.22, -0.81** | **0.001** |
| RMR | 0.47 | 0.41 | 1.60 | -0.26, 1.31 | 0.26 |
| **$M_{\text{sum}}$** | **-1.04** | **0.45** | **0.35** | **-1.93, -0.15** | **0.02** |
| **Endurance × $M_{\text{sum}}$** | **-2.14** | **0.95** | **0.12** | **-4.00, -0.29** | **0.02** |
| Conductance | 0.09 | 0.24 | 1.09 | -0.41, 0.98 | 0.70 |
| Tarsus | -0.06 | 0.19 | 1.06 | -0.85, 0.37 | 0.75 |
| $M_b$ | 0.06 | 0.23 | 1.06 | -0.57, 1.02 | 0.80 |

## DISCUSSION

To support their energetic lifestyle, homeothermic endotherms maintain a relatively high and constant T_b_ despite changes in the environment. Regulating T_b_ within this narrow window necessitates responding to changes in their environment that may arise both predictably and stochastically. Here we show that the capacity for T_b_ maintenance is a flexible avian phenotype that can acclimate to changes in the thermal environment. The ability to maintain normothermia during acute cold exposure improved with cold acclimation, as well as the duration of the acclimation treatment. Modifications to thermoregulatory ability occurred on relatively short time scales (within one week) and without changes in photoperiod, suggesting that juncos can match their thermoregulatory physiology to current thermal conditions independent of broad-scale seasonal cues. At the same time, further enhancements to the ability to maintain T_b_ were made over successive time steps, indicating a lag in the induction of some physiological modifications. These results emphasize the potential for temporal constraints on individual flexibility.

### Correlates of improved T_b_ maintenance ability

Summit metabolic rate has previously been implicated as the main factor governing avian cold tolerance in studies of seasonal flexibility (Swanson, 2010). We found that M_sum_ increased with cold acclimation within one week of cold exposure, but that further enhancements to this trait did not occur with longer acclimation durations. In this respect, our study is unique in that it shows responses in M_sum_ occurring on the order of days rather than weeks or months. Furthermore, our results indicate that the magnitude of the change in M_sum_ over this short timescale is on the order of seasonal increases in M_sum_ exhibited in wild juncos between summer and winter (28%, Swanson, 1990a), as well as that previously shown for juncos exposed to laboratory acclimations under more moderate conditions (16-19% at 3°C for six weeks, Swanson et al., 2014). The comparable magnitude of response to these two different temperature treatments contrasts with previous work showing that wild juncos and other birds modulate M_sum_ with environmental temperature across the winter (Swanson & Olmstead, 1999). Taken together, these findings suggest that M_sum_ might be coarsely adjusted, rather than fine-tuned, to environmental temperature, and that there may be limits to their flexibility in response to temperature variation (Petit and Vézina, 2014). Dissecting the relative contribution of subordinate phenotypic traits to M_sum_— *e.g.,* pectoralis muscle size, hematocrit, or cellular metabolic intensity (Liknes & Swanson, 2011; Swanson, 1990b; Swanson et al., 2014)— will illustrate how birds build this phenotype and which traits (if any) may be limiting its flexibility.

Individuals characterized by both elevated M_sum_ and the ability to sustain heightened M_sum_ (endurance) were also capable of maintaining normothermia longer, indicating an additive effect of enhancing these two phenotypes. Nonetheless, we saw no effect of acclimation treatment or duration on endurance and individuals continued to enhance their ability to maintain normothermia in successive weeks long after M_sum_ plateaued. These results suggest that either these indices are insufficient indicators of total thermogenic capacity or that individuals reduced their thermal conductance at these later time points.

In support of this latter possibility, we found that conductance of the skin and plumage decreased in response to our temperature stimulus. This finding prompts questions about the exact mechanism underlying such a modification. Although we cannot distinguish between potential adjustments made to the properties of the skin or the plumage, the fact that heat loss was only reduced at the last sampling point (*Week 9*) suggests that alterations to thermal conductance may require significant time to implement. We did not see evidence that birds were molting large amount of feathers during the acclimation, as was obvious when birds first entered captivity (Stager *pers. obs.*). Moreover, avian molt is closely tied to photoperiod (Danner et al., 2015), yet conductance changed in the absence of variation in photoperiod. Instead, it seems plausible that birds may have added body feathers to their existing plumage. Previous work has shown that juncos increase plumage mass in winter compared to summer (Swanson, 1991), as do American Goldfinches (*Carduelis tristis*), which additionally have been shown to possess a greater percentage of plumulaceous barbules, as well as more barbules per barb, in winter than summer (Middleton, 1986). However, goldfinches undergo an alternate molt in the spring, in addition to the basic molt in autumn, whereas juncos exhibit just the single autumn molt (Pyle, 1997). Thus if juncos did selectively add feathers to reduce conductance in the cold, it would suggest that they concomitantly lose select feathers before the subsequent summer to enable increased heat loss when they need it most. Alternatively, it is possible that changes were made to the heat transfer properties of the skin itself. For example, avian skin composition can be flexibly remodeled on the time scales of our experiments in response to humidity (Muñoz‐Garcia et al., 2008). It should be noted that while the *Week 9* treatment was our smallest sample size, our results are statistically robust. Future studies would therefore profitably combine our methodology here with data on the time course of plumage quality and mass to further elucidate the role that heat saving mechanisms might play in avian T_b_ maintenance.

While reduced thermal conductance may explain the final boost in ability to maintain normothermia seen at *Week 9*, variation in neither M_sum_ or conductance explain the increase in T_b_ maintenance at *Weeks 2* and *6*. One potential reason for this disparity is that we were unable to quantify total heat loss in live birds and thus may have overlooked additional factors that contribute to minimum conductance — like vasoconstriction (Irving and Krog, 1955), posture (Pavlovic et al., 2019), and ptiloerection (Hohtola, Rintamäki, & Hissa, 1980) — that may have varied across treatments. To this point, we can anecdotally report from observations made during cold exposure trials that juncos sat on their feet, puffed up their feathers, but did not tuck their heads under their wings; however, we did not quantify these postures. A second potential explanation is that assaying total oxygen consumption could mask potential changes to thermogenic efficiency. For example, juncos may achieve higher metabolic efficiency by increasing fiber size within their muscle, thereby allowing for greater contraction force while simultaneously reducing basal metabolic cost because larger muscle fibers require less energy by Na^+^/K^+^ ATPase to maintain sarcolemmal membrane potential (Jimenez et al., 2013). Such changes have been documented in Black-capped chickadees (*Poecile atricapillus*), which exhibit seasonal decreases in muscle fiber diameter from spring to summer (Jimenez et al., 2019), as well as increases with cold acclimation (Vezina et al., 2020). Additionally, if adult birds are employing non-shivering thermogenesis, the relative proportion of shivering to non-shivering processes could be altered seasonally. Direct measures of shivering and/or non-shivering thermogenesis, however, are needed to test for these potential changes. Our results thus point to exciting directions for further exploration regarding the mechanisms governing seasonal acclimatization in avian T_b_ maintenance.

### Thermoregulation and broad-scale ecogeographic patterns

Spatial variation in basal metabolic rate (BMR) is often interpreted as a thermal adaptation to cold conditions, whereby colder climates are correlated with higher endothermic BMR (Lovegrove, 2003; Wiersma et al., 2007). Changes in BMR have also been implicated as a mechanism and/or by-product of avian thermal acclimation across seasons (Dutenhoffer and Swanson, 1996). Here we did not find increases in RMR associated with cold acclimation. We quantified RMR rather than BMR, meaning that birds were not fasted before measurements. Nonetheless, RMR post-acclimation was similar to previously published BMR values for wild juncos (Swanson et al., 2012). We found that RMR was not correlated with other performance phenotypes (M_sum_, conductance, or T_b_ maintenance), implying that it is not a good indicator of avian cold tolerance. This result also agrees with previous work showing that M_sum_ and RMR can be uncoupled (Petit et al., 2013; Swanson et al., 2012). Finally, it indicates that the energetic costs associated with enhancing thermoregulatory ability — like building the metabolic machinery associated with increased M_sum_ — do not necessarily manifest as higher resting energetic use.

M_sum_ is commonly used as a proxy for cold tolerance in macrophysiological studies (*e.g.,* Stager et al., 2016). However, our results highlight a disconnect between these two measures. Although junco M_sum_ was correlated with T_b_ maintenance in the cold, it was not as strong a predictor of T_b_ maintenance as was endurance, and it was the interaction between M_sum_ and endurance that had the largest effect on T_b_ maintenance. Furthermore, the amount of variation in T_b_ maintenance explained by M_sum_ alone was relatively small. These results echo those of a previous study in which variation in M_sum_ did not match variation in cold tolerance in two other, disparate, junco populations (Swanson, 1993). M_sum_ may, therefore, not be as strong a proxy of cold tolerance as frequently thought. Nonetheless, to discern whether this pattern can be generalized to other taxa, we encourage the collection of T_b_ data to assess normothermic ability as we have done here. Such data is increasingly easy to obtain using PIT tags and other next-generation tracking technologies (*e.g.*, Parr et al., 2019)

### Responding to fluctuating environmental conditions

Nicknamed “snowbirds” for their winter tenacity, juncos are not unique in their cold hardiness. Their close relative, the White-throated Sparrow (*Zonotrichia albicolli*), has been acclimated to even colder conditions than those employed here (3 wk at −20° C) (McWilliams and Karasov, 2014), and other small songbirds have survived short periods in the lab at −60° C (Dawson and Carey, 1976). Given that the climatic conditions juncos experience vary across their broad geographic distribution, junco populations may also differ in their thermoregulatory abilities and the underlying physiological responses they use to moderate T_b_. Acclimatizing to these cold temperatures in the wild likely comes with tradeoffs, such as increased exposure to predators as a consequence of increased time spent foraging (Lima, 1985). Moreover, as our results demonstrate, the duration of the cold period may dictate which physiological strategies are utilized. For instance, we found that juncos are capable of responding to thermal cues with large changes in M_sum_ occurring within one week. However, rapid changes likely require energetic input to fuel this physiological remodeling, in addition to those required to elevate aerobically powered shivering thermogenesis.

Another short-term strategy that birds use to cope with cold temperatures is facultative heterothermia (Mckechnie and Lovegrove, 2002). We witnessed similar patterns of oscillating T_b_ in some juncos, whereby they raised T_b_ to normothermic levels following a period of hypothermia. Counter to previous findings (Swanson, 1991), this suggests that juncos may employ facultative heterothermia as an energy saving mechanism. However, we did not find evidence for acclimation in this strategy — as members of both temperature treatments exhibited heterothermia — nor that birds differed in their starting T_b_ among temperature treatments. The White-crowned Sparrow (*Z. leucophrys*), another close relative of the junco, has been shown to lower their T_b_ by 3.6°C (Ketterson and King, 1977), but we found that juncos could lower their T_b_ by as much as 7°C and still recover normothermia during an acute cold trial. Although we did not assess potential consequences of hypothermia in this context, 7°C is well within the range of T_b_ reductions observed in other passerines (Mckechnie and Lovegrove, 2002). Furthermore, a nightly reduction in T_b_ of this magnitude is estimated to reduce the energy expenditure of *Parus* tits by up to 30% and increase their over-wintering survival by 58% (Brodin et al., 2017). Like other birds, however, juncos suffer impaired mobility at such low T_b_ (Stager *pers. obs.*). While rest-phase hypothermia may be especially useful at night when activity levels are reduced, it alone may not be a good strategy to cope with cold temperatures during the day when birds need to eat, move, and avoid predators (Brodin et al., 2017).

Juncos may thus be layering longer-term modifications — like the observed changes in conductance — on top of these shorter-term mechanisms to arrive at the optimal phenotype for the challenge at hand. If widespread, this would provide birds with a host of strategies to employ, each of which may be useful over different time scales. As a result, in the face of increasing climatic variability, some birds may be well equipped to deal with potential mismatches between photoperiod and temperature that lead to thermoregulatory challenges in the cold. However, their ability to employ these different strategies is likely dependent on their access to sufficient food to fuel and maintain these phenotypic changes. Because food resources are also likely to vary in response to global change (Rafferty, 2017; Williams and Jackson, 2007), future work should investigate the complex interactions between environmental change, subsequent physiological responses, and their energetic costs.

## ACKNOWLEDGEMENTS

We are especially thankful to Phred Benham, Ryan Mahar, Nick Sly, Hannah Specht, and Stan Senner for their assistance in catching birds. We also thank Frank Moss for logistical assistance, Hailey Bunker for help with husbandry, and Keely Corder, Rena Schweizer, Jon Velotta, and Cole Wolf for comments on an earlier version of this manuscript.

## AUTHORS’ CONTRIBUTIONS

MS and ZAC conceived of the study; NRS helped capture birds; BWT designed the conductance trials; MS performed all data collection and analyses, and drafted the manuscript; NRS, BWT, and ZAC contributed edits to the manuscript. All authors gave final approval for publication.

## DATA AVAILABILITY

All data are available in the electronic supplementary material.

## ETHICS

All procedures were approved by the University of Montana Animal Care Committee (Protocol 010-16ZCDBS-020916). Birds were collected with permission from Montana Fish Wildlife & Parks (permits 2016-013 and 2017-067-W, issued to MS) and the US Fish & Wildlife Service (permit MB84376B-1 to MS).

