## Supplemental Tables and Figures for "Body temperature maintenance acclimates in a winter-tenacious songbird"

**Supplementary Materials**

**Table S1.** Sex distribution of treatment groups.

|  | **Week 1** | | **Week 2** | | **Week 3** | | **Week 6** | | **Week 9** | |
| --- | --- | --- | --- | --- | --- | --- | --- | --- | --- | --- |
| **Cold** | 11 | 1 | 10 | 2 | 11 | 1 | 10 | 2 | 5 | 0 |
| **Control** | 10 | 1 | 11 | 1 | 11 | 1 | 11 | 1 | 6 | 0 |
|  | M | F | M | F | M | F | M | F | M | F |

**Table S2.** Effect of respirometry unit on pre-acclimation RMR in (a) 2016 and (b) 2017 and on M_sum_ in (c) 2016 and (d) 2017. Unit A is reference for 2016 and Unit 3 is reference for 2017.

a.

| **Unit** | **β** | **SE** | **p** |
| --- | --- | --- | --- |
| Unit B | -0.07 | 0.06 | 0.24 |
| Unit C | 0.12 | 0.05 | 0.04 |

b.

| **Unit** | **β** | **SE** | **p** |
| --- | --- | --- | --- |
| Unit 1 | 0.13 | 0.11 | 0.21 |
| Unit 4 | -0.52 | 0.06 | 2.4 x 10^-10^ |

c.

| **Unit** | **β** | **SE** | **p** |
| --- | --- | --- | --- |
| Unit C | -0.17 | 0.15 | 0.25 |

d.

| **Unit** | **β** | **SE** | **p** |
| --- | --- | --- | --- |
| Unit 1 | 7.94 | 0.58 | < 2 x 10^-16^ |
| Unit 4 | -1.54 | 0.25 | 2.5 x 10^-7^ |
| Unit 5 | 4.62 | 0.34 | < 2 x 10^-16^ |

**Table S3.** Effect of thermal conductance assay date on average power input.

| **Date** | **β** | **SE** | **p** |
| --- | --- | --- | --- |
| 5/20/19 | -1.24 | 17.21 | 0.94 |
| 5/21/19 | 21.89 | 16.39 | 0.19 |
| 5/22/19 | 4.57 | 17.21 | 0.80 |
| 5/23/19 | 13.39 | 16.76 | 0.43 |
| 5/24/19 | - 1.01 | 20.84 | 0.96 |
| 5/27/19 | 9.40 | 20.84 | 0.65 |
| 6/6/19 | 26.99 | 18.50 | 0.37 |
| 6/7/19 | 21.50 | 16.76 | 0.46 |
| 6/8/19 | -16.84 | 17.21 | 0.48 |
| 6/9/19 | 12.37 | 17.21 | 0.75 |
| 6/10/19 | -12.25 | 17.21 | 0.12 |
| 6/11/19 | -5.2 | 16.76 | 0.20 |

**Table S4.** Survival model output for hypothermic state (defined as < 37°C) as a function of cabinet temperature, acclimation temperature treatment, and duration treatment. Negative β coefficients represent reduced risk of hypothermia. Hazards ratio (HR) is the exponent of the β coefficient (*i.e.* a reduction in the hazard by this factor). *Control* treatment is reference for Temperature effect. All continuous variables were standardized; bold indicates predictor variables with statistically significant effects on T_b_ maintenance.

| Variable | *β* | SE | HR | 95% CI | *p* |
| --- | --- | --- | --- | --- | --- |
| Cabinet | **-3.70** | **0.49** | **0.02** | **-4.98, -2.41** | **1.7 x 10^-8^** |
| Treatment | **-2.00** | **0.41** | **0.14** | **-3.13, -0.87** | **5.2 x 10^-4^** |
| Period | **-1.30** | **0.24** | **0.27** | **-1.99, -0.60** | **2.4 x 10^-4^** |

**Table S5.** Highest-ranked models (with lowest AIC_c_ scores) in candidate set for effects of phenotypic variables on the maintenance of T_b_ when hypothermic state was defined as < 37°C using Cox proportional hazards models. Only models with ΔAIC_c_ < 4 are reported. *K* indicates the number of parameters in each model; Cabinet refers to the cabinet temperature during the cold trial.

| Candidate Model | *K* | AIC_c_ | ΔAIC_c_ | *w_i_* |
| --- | --- | --- | --- | --- |
| Cabinet + Conductance + Endurance × M_sum_ + RMR | 6 | 989.3 | 0.0 | 0.13 |
| Cabinet + Conduct. + M_b_ + Tarsus + Endurance × M_sum_ + RMR | 8 | 989.3 | 0.0 | 0.13 |
| Cabinet + Conductance + Endurance × M_sum_ + RMR + Tarsus | 7 | 989.5 | 0.3 | 0.11 |
| Cabinet + Endurance × M_sum_ + M_b_ + RMR + Tarsus | 7 | 990.1 | 0.8 | 0.09 |
| Cabinet + Conductance + Endurance x M_sum_ + M_b_ +RMR | 7 | 990.2 | 1.0 | 0.08 |
| Cabinet + Endurance × M_sum_ + RMR | 5 | 990.4 | 1.2 | 0.07 |
| Cabinet + Endurance × M_sum_ + RMR +Tarsus | 6 | 990.4 | 1.2 | 0.07 |
| Cabinet + Endurance × M_sum_ + M_b_ + RMR | 6 | 991.2 | 2.0 | 0.05 |
| Cabinet + Conductance + Endurance + M_sum_ + RMR | 5 | 992.1 | 2.8 | 0.03 |
| Cabinet + Conductance + Endurance × M_sum_ + M_b_ | 6 | 992.2 | 2.9 | 0.03 |
| Cabinet + Endurance + RMR + M_sum_ | 4 | 992.5 | 3.2 | 0.03 |
| Cabinet + Conductance + Endurance × M_sum_ | 5 | 992.6 | 3.4 | 0.02 |
| Cabinet + Conductance + Endurance × M_sum_ + M_b_ + Tarsus | 7 | 992.8 | 3.6 | 0.02 |
| Cabinet + Conductance + Endurance + M_b_ + RMR + M_sum_ | 6 | 993.0 | 3.7 | 0.02 |

**Table S6.** Linear effects of *Cold* treatment, *Duration,* and their interaction on phenotypic traits before and after acclimation. Mass (M_b_) is included as a covariate for metabolic traits. Delta (Δ) represents change over acclimation period (post- minus pre-acclimation) for traits that were measured at both time points. Metabolic rates are expressed as measures of oxygen consumption per (ml O_2_^1^•min^-1^); conductance expressed in mW; endurance in min; mass in g. Bolded significant effects after Bonferroni correction (p < 0.004). Sample sizes reported in Table 1.

| **Phenotype** | | **Intercept** | | **M_b_** | | | **Cold Treatment** | | | **Duration** | | | **Treat x Duration** | | |
| --- | --- | --- | --- | --- | --- | --- | --- | --- | --- | --- | --- | --- | --- | --- | --- |
|  |  | **β** | **SE** | **β** | **SE** | **p** | **β** | **SE** | **p** | **β** | **SE** | **p** | **β** | **SE** | **p** |
| Pre | M_b_ | 22.51 | 0.38 |  |  |  | -0.32 | 0.54 | 0.55 | -0.10 | 0.08 | 0.26 | 0.10 | 0.12 | 0.43 |
|  | Tarsus | 19.92 | 0.14 |  |  |  | 0.07 | 0.20 | 0.73 | 0.04 | 0.03 | 0.25 | 0.01 | 0.04 | 0.79 |
|  | RMR | 0.22 | 0.32 | **0.05** | **0.01** | **7.2 x 10^-4^** | 0.05 | 0.08 | 0.49 | 0.00 | 0.01 | 0.68 | -0.02 | 0.02 | 0.32 |
|  | M_sum_ | 4.77 | 0.80 | 0.07 | 0.04 | 0.06 | 0.30 | 0.21 | 0.16 | 0.01 | 0.03 | 0.76 | -0.05 | 0.05 | 0.25 |
|  | Endur. | 39.39 | 18.41 | -0.50 | 0.81 | 0.54 | -1.49 | 4.91 | 0.76 | -0.33 | 0.75 | 0.66 | 0.81 | 1.09 | 0.46 |
| Post | M_b_ | 23.08 | 0.40 |  |  |  | -0.03 | 0.56 | 0.95 | -0.20 | 0.09 | 0.03 | 0.14 | 0.13 | 0.27 |
|  | RMR | 0.40 | 0.38 | **0.05** | **0.02** | **3.6 x 10^-3^** | -0.14 | 0.09 | 0.15 | -0.03 | 0.02 | 0.05 | 0.03 | 0.02 | 0.19 |
|  | M_sum_ | 5.09 | 1.76 | 0.08 | 0.07 | 0.31 | **1.35** | **0.44** | **2.9 x 10^-3^** | -0.09 | 0.07 | 0.20 | -0.01 | 0.10 | 0.93 |
|  | Endur. | 16.50 | 22.44 | 0.19 | 0.94 | 0.85 | -4.66 | 5.62 | 0.41 | 1.43 | 0.91 | 0.12 | -0.37 | 1.26 | 0.77 |
|  | Conduct. | 308.62 | 7.96 |  |  |  | 30.10 | 11.28 | 9.0 x 10^-3^ | 3.06 | 1.78 | 0.09 | **-8.52** | **2.56** | **1.2 x 10^-3^** |
| Δ | M_b_ | 0.57 | 0.46 |  |  |  | 0.29 | 0.65 | 0.66 | -0.10 | 0.10 | 0.32 | 0.04 | 0.15 | 0.76 |
|  | RMR | 1.52 | 0.07 | 0.02 | 0.01 | 0.11 | -0.15 | 0.10 | 0.14 | -0.04 | 0.02 | 0.02 | 0.03 | 0.02 | 0.12 |
|  | M_sum_ | 6.92 | 0.31 | -0.11 | 0.07 | 0.09 | **1.32** | **0.43** | **3.0 x 10^-3^** | -0.12 | 0.07 | 0.07 | 0.01 | 0.10 | 0.94 |

**Table S7.** Excel file with raw data: ID, Treatment, Duration, Year, Sex, Tarsus, Masses, RMR, M_sum_, Endurance, Conductance, Time to Hypothermia for each individual.

**Figure S1.** Minimum temperature for North America using WorldClim data and winter junco distribution demarcated with white lines (approximated from Nolan et al., 2002).

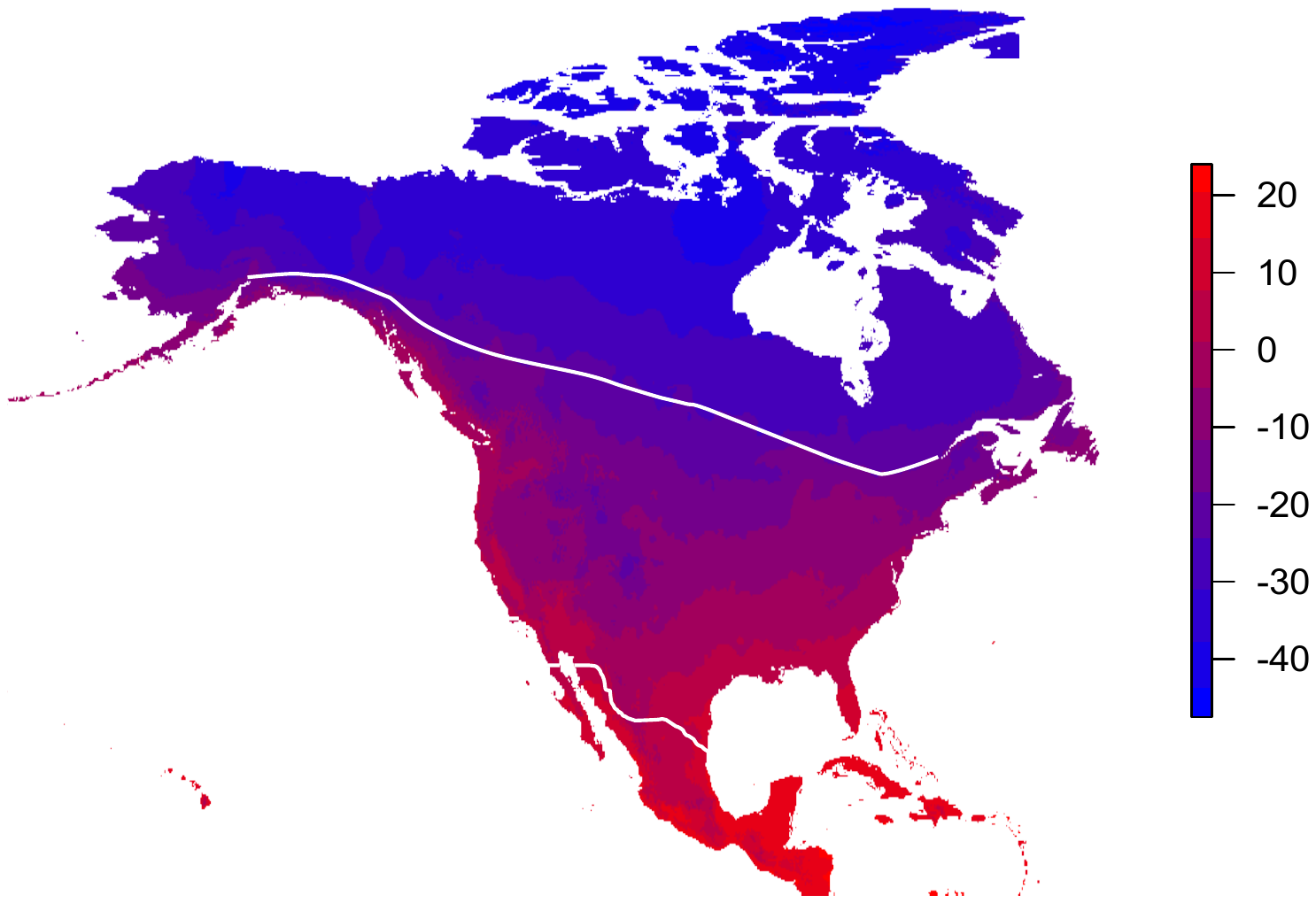
